# Maintenance of cooperation in a yeast population in a public-good driven system

**DOI:** 10.1101/2023.06.01.543253

**Authors:** Namratha Raj, Supreet Saini

**Affiliations:** Department of Chemical Engineering, Indian Institute of Technology Bombay, Mumbai 400076

**Keywords:** Cooperation, cheating, public good system, microbe, SNV/Indel, CNV

## Abstract

The phenomenon of cooperation is prevalent at all levels of life. In microbial communities, some groups of cells exhibit cooperative behaviour by producing costly extracellular resources that are freely available to others. These resources are referred to as public goods. *Saccharomyces cerevis*iae secretes invertase (public good) in the periplasm to hydrolyse sucrose into glucose and fructose, which are further imported by the cells. After hydrolysis of sucrose, the cells retain only 1% of the monosaccharides, while 99% diffuse into the environment and can be utilised by all neighbouring cells. The non-producers of invertase (cheaters) exploit the invertase-producing cells (cooperators) by utilising the monosaccharides and paying nothing for the latter. In this work, we investigate the evolutionary dynamics of this cheater-cooperator system. If cheaters are selected for their higher fitness, the population will collapse. For cooperators to survive cheating and thrive in nature, they should have evolved some survival strategies. To understand the adaptation of cooperators in sucrose, we performed a coevolution experiment in sucrose. Our results show that cooperators increase in fitness as the experiment progresses. This phenomenon was not observed in environments which involved a non-public good system. Genome sequencing reveals the molecular basis for the cooperator adaptating is because of increased privatization of the public-good released carbon resource.

## Introduction

Microorganisms communicate and cooperate with each other to perform various activities, such as nutrient acquisition, biofilm formation, among other functions(1-4). Often, members of a microbial population produce extracellular resources, such as an enzyme or a metabolite, to achieve these functions(5, 6). Since these products are released in the extracellular environment and both producers and non-producers enjoy the benefits of these secretions; these extracellular secretions are called public goods. The microbes that produce these public goods by bearing the costs of production are called cooperators, while cheaters consume public goods and do not contribute towards their production(7, 8).

This suggests that, in general, cheaters will have a fitness advantage over cooperators and are thus, expected to expand in the population. Such a scenario suggests that the population could be vulnerable to a crash, especially in a case where the public good is essential for growth and/or survival(9, 10).

In such a setting, how does cooperation, as a trait, survive?

Cooperators have evolved many mechanisms to protect themselves from cheaters. Production of public goods in bacterial communities is mainly regulated by quorum sensing(11, 12). The microbes produce extracellular chemicals called autoinducers and when the concentration of autoinducers detected by the cells exceeds a minimum threshold, the cells begin to express the genes to produce public goods(11). Cooperators also target benefits and punishment as mechanisms to protect themselves from exploitation by cheaters(13, 14). For example, in *Dictyostelium discoideum*, a low sporulation efficiency was observed in cheaters (which do not contribute to the formation of multicellular fruiting bodies under starvation)(15). In targeted punishment, cheaters are punished by cooperators for cheating. The quorum cascade system of *Pseudomonas aeruginosa* has one such mechanism. Its LasR-LasI quorum sensing system regulates the production of many public goods such as proteases and virulence factors(16). The LasR-LasI quorum sensing system also controls another RhII-RhIR quorum system that produces cyanide, a toxin to punish cheaters. Cooperators, on the other hand, are less susceptible to cyanide(13). Lastly, cheaters outgrowing cooperators is often frequency dependent. Partial privatisation of the public good by the cooperators also prevents cheaters from driving cooperators to extinction(10, 17, 18).

Little is known about the strategies that yeast utilizes to prevent cheating in public good systems. Adaptive race model was proposed to support the prevalence of cooperators in the public good system(9). In order to study cheater-control mechanisms, Waite and colleagues started a coevolution experiment with equal ratios of three engineered strains (two types of cooperators and one cheater). The authors observe that often the population crashed, due to the cheater outcompeting the cooperators. Many theoretical studies have been carried out based on the experimental results of cheater and cooperator coevolution to understand their adaptation. Game theory has been extensively used to study the cheater-cooperator interaction and evolution(17, 19) and cooperator behaviour has been explained using prisoner’s dilemma, snowdrift games(17, 19). The analysis of single cell cost-benefit dynamics predicts that cooperators with ability to retain optimal fraction of public goods can perform better than cheaters(20). The findings suggest that cooperators might find a balance between cooperative behaviour and survival by partial privatization. In the face of the appearance of cheaters or ecological stress factors, it is predicted that the evolution of facultative cooperative behaviour will increase the cooperator survival(21).

Yeast cannot import disaccharides sucrose, melibiose or a trisaccharide like raffinose. Instead, they secrete enzymes for the hydrolysis of di-/trisaccharide either in periplasmic or extracellular regions and the resulting monosaccharides are then transported into the cells(22, 23). *S. cerevisiae* hydrolyses sucrose into glucose and fructose in the periplasm, using an invertase encoded by the gene SUC2(24-27). Roughly ∼99% of the resulting monosaccharides are released into the media making them available for utilization by any cell; while the small remaining fraction is retained by the cell producing the enzyme (public good)(17) (**Figure 1**).

**Figure 1.**
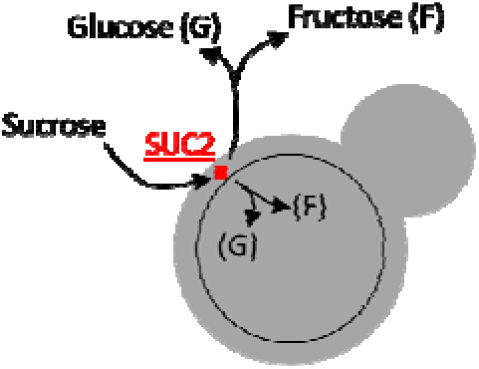
Sucrose utilization in *S. cerevisiae* is facilitated by invertase SUC2p, which is secreted in the periplasm, where it hydrolyses sucrose into glucose and fructose. Roughly 99% of the hydrolysed monosaccharides are released into the media, and are available to all members of the population. The remaining amount is retained by the cell that produces SUC2p.

In a non-repressing medium, SUC2 is expressed at a basal level. It is induced eight times more than basal level when there is a small amount of glucose or fructose present in the medium (<0.1% w/v)(28). Following hydrolysis of sucrose, small amounts of glucose and fructose induce SUC2 expression by phosphorylating SUC2 repressors Mig1, Mig2, and Rgt1 by the Snf1/Snf4 complex(29). On the other hand, in response to high glucose levels in the environment Mig1 and Mig232 inhibit SUC2 transcription(30).

To test adaptation of cheaters and cooperators, when grown together, we perform a coevolution experiment with a cooperator and cheater strain of yeast, using the sucrose utilization system. In the environment of choice, cheaters have a fitness advantage over the cooperators. However, after coevolution for 200 generations, we show that the cooperators adapt to perform better than the cheaters, and hence, increase in frequency. This behaviour was observed in all six independent lines of the experiment, and was contingent on a public good system. Genome sequencing reveals the targets responsible for this adaptive behaviour.

## Results

### Sucrose utilization is contingent on strains carrying SUC2

Growth kinetics of wild type and the ΔSUC2 mutant in an environment containing sucrose as the carbon source was studied. The wildtype BY4741 (cooperator) and the ΔSUC2 (cheater) exhibit qualitatively different growth kinetics in this environment (**Figure 2**). While the cooperator exhibits growth in this environment, the cheater is unable to grow in the sucrose environment.

**Figure 2.**
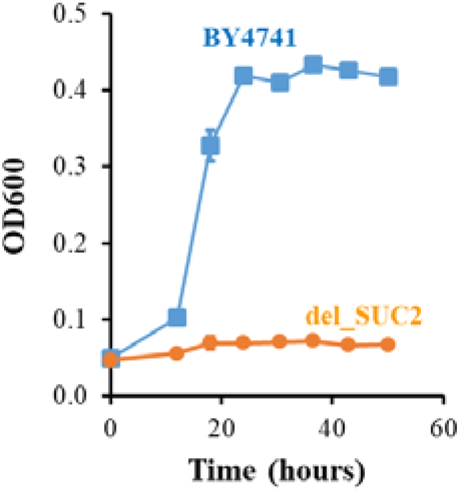
The BY4741 (wildtype/cooperator) strain having SUC2 gene grows in 0.2% sucrose media, while BY4741 derived ΔSUC2 (cheater) did not grow in sucrose. The experiment was performed in triplicate, and the mean and the standard deviation are reported. The growth rates are significantly different with p-value<0.001, unpaired t-test.

As ΔSUC2 is unable to grow in sucrose, we coevolved strain with wild-type BY4741 for 200 generations. In the experiment, wildtype was the cooperator strain, and ΔSUC2 was the cheater strain. The cheater strain was dependent on wild type for release of glucose and fructose, and hence growth. As control, these two strains were also coevolved in environments containing (a) glucose, (b) fructose, and (c) mixture of glucose and fructose.

### In a batch culture, cooperators have lower fitness than cheaters

Six independent coevolution lines of cheaters and cooperators were initiated in media containing 0.2% sucrose. The culture was started with 1:1 ratio of the two genotypes (BY4741 and ΔSUC2). To quantify the cost associated with cooperation, the frequency of cooperators and cheaters were calculated after a day of growth. Our results show that the ratio of the frequency of cooperators to cheaters was 3:7 (**Figure 3A**). This implied that ΔSUC2 (cheaters) are more fit than wildtype in the sucrose environment.

**Figure 3.**
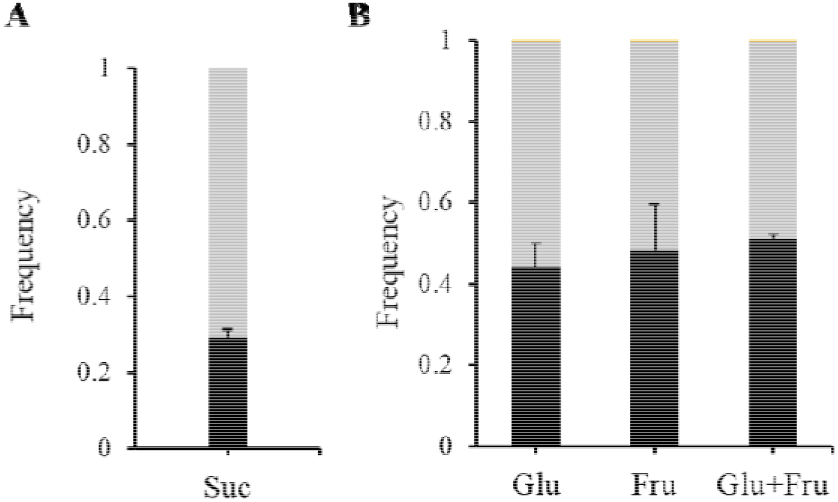
**(A)** The cooperator BY4741 and cheater ΔSUC2 were grown in 0.2% sucrose. After a day of growth, the frequencies of both strains were quantified. The ratio of initial frequency of cooperators to frequency of cheaters in all six experimental lines was approximately 3:7. Cooperator frequency is in black; cheater frequency is in grey. The average and standard deviation of the six lines is shown in the Figure. **(B)** The BY4741 and ΔSUC2 were grown in (a) 0.2% glucose (Glu), (b) 0.2% fructose (Fru), and (c) 0.1% glucose+0.1% fructose (Glu+Fru). After a day of growth, the frequencies of both strains were quantified in all the three replicate lines of each media. The ratio of initial frequency of cooperators to frequency of cheaters was around 1:1 in all the lines of all the media. Cooperator frequency is in black; cheater frequency is in grey. All experiments were performed in triplicate. The average and standard deviation are reported.

To determine if the fitness cost of the wildtype was due to cooperative invertase production, identical competition experiments were conducted in three different media containing glucose, fructose, and mixtures of both sugars. The initial ratio of frequency of cooperators to cheaters was 1:1 in all these non-public good systems (**Figure 3B**). The results showed that both cooperators and cheaters were present in an equal frequency in a system where a public good was not required for growth. This demonstrates that in the presence of cheaters in sucrose, cooperators suffered from the cost of production of invertase.

### Cooperator, upon evolution in sucrose, adapts to perform better than cheaters

Cooperators and cheaters were coevolved via a 1:100 dilution after every 24 h of growth in 0.2% sucrose. A single dilution of 1:100 corresponds to approximately 6.7 generations and the coevolution experiment was continued for 200 generations. Over the course of the experiment, the frequency of the cooperators increased in all six experimental lines (**Figure 4**).

**Figure 4.**
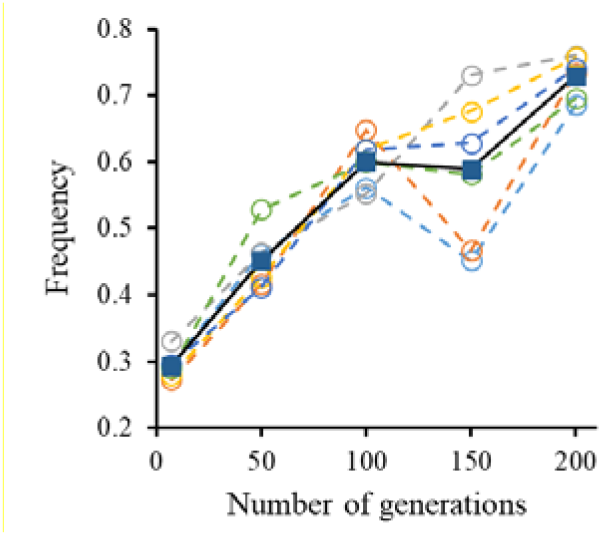
The coevolution of cooperator BY4741 and cheater ΔSUC2 in 0.2% sucrose was started with 1:1 ratio of both genotypes. The frequency of cooperators and cheaters were quantified every 50 generations. The frequency of cooperators in the co-culture significantly increased in the 200 generations (p-value<10^−10^, unpaired t-test). Dotted line represents individual lines. Solid line represents the average of the six lines.

After 50th generation, the population’s frequency of cooperators began to rise after initially being low. Studies on the coevolution of cheaters and cooperators in sucrose have revealed that frequency/density dependent selection may cause frequency of cooperators to rise until they reach a steady state of coexistence with cheaters after which the cheaters might increase in frequency again(10, 17, 31, 32). However, neither a steady state nor a rise in the frequency of cheaters was observed in our experiment even after the 100th generation. At 200th generation of coevolution the ratio of frequency of cooperators to cheaters was 7:3; this increase in frequency is statistically significant (p-value<10^−10^, unpaired t-test) (**Figure 4**).

### Increase in fitness of cooperators is not facilitated by increase in growth rate

After coevolution for 200 generations, the frequency of cooperators was significantly more than that of the cheaters. However, somewhat surprisingly, the growth kinetics of the evolved cooperator was found to be qualitatively similar to the ancestors (**Figure 5**).

**Figure 5.**
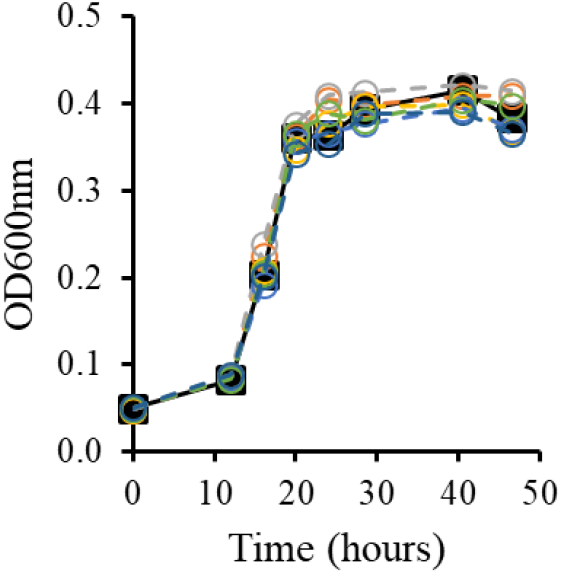
Growth kinetics of ancestor cooperator (solid) and the coevolved cooperators at 200^th^ generation of all six lines (dashed) are represented. The growth rate of cooperators of lines are not statistically different from that of ancestor (p-value >0.5, unpaired t-test).

To determine if the coevolved cooperators perform better against the ancestor cheater, a competition assay was conducted with ancestor cheater and coevolved cooperators. The frequency of coevolved cooperators of each line was higher than ancestor cheater after 24 h of growth in 0.2% sucrose (**Figure 6A**). These results are significantly different than the initial observation obtained before in the competition between ancestor cooperator and ancestor cheater (p-value<10^−6^, unpaired t-test) (Figure 3A). Coevolved cooperators at the 200th generation were outperforming ancestor cooperators in four lines despite no significant increase in growth rates. These results show that the cooperators had not increased their growth rate but they had employed some other mechanism to proliferate in the population.

**Figure 6.**
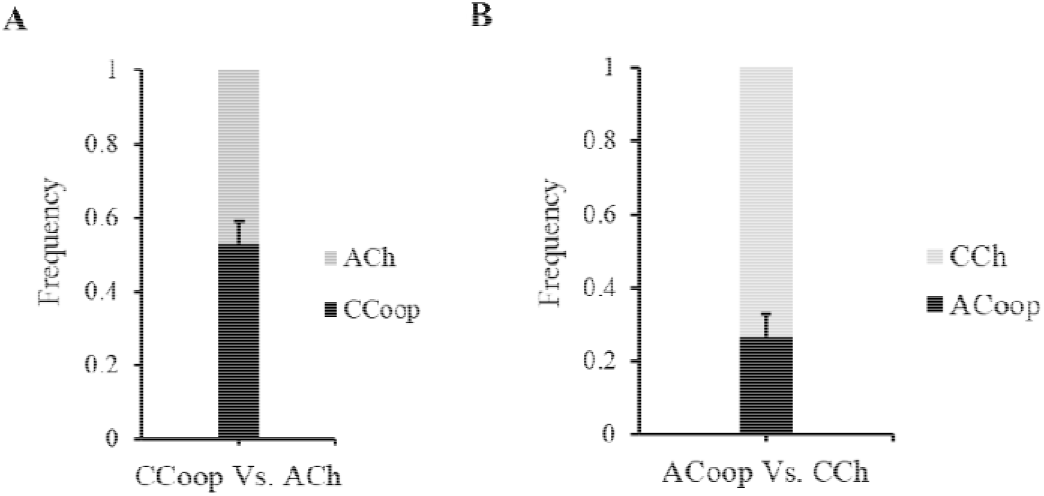
**(A)** At 200^th^ generation, how did coevolved cooperator (CCoop) perform against ancestor cheater (Ach)? To understand this, competition between evolved cooperators (of all six lines) and ancestor cheater was conducted. The fraction of cooperators was more than 50%. These results are significantly different than the initial observation obtained from the competition between ancestor cooperator and ancestor cheater (p-value<10^−6^, unpaired t-test) (Figure 3A). **(B)** At 200^th^ generation, to understand how coevolved cheaters (CCh) at performed against ancestor cooperator (ACoop) the competition assay between coevolved cheaters (of all six lines) and ancestor cooperator was conducted. The fraction of cheaters was around 70%. These results are not significantly different than the initial observation obtained from the competition between ancestor cooperator and ancestor cheater (p-value ∼ 0.35, unpaired t-test).

Also, the ratio of frequency of cheaters to cooperators was almost 7:3 when ancestor cooperator competed with evolved cheaters. These results are qualitatively similar to the initial observation obtained in the competition between ancestor cooperator and ancestor cheater (p-value ∼ 0.35, unpaired t-test) (**Figure 6B**).

### Whether the adaptation of BY4741 to do better than ΔSUC2 is public-good driven? How do cooperators evolve in non-public goods systems?

We observed how cooperators evolved to perform better than cheaters in sucrose which is a public good system. Since both the strains utilize monosaccharides equally well, neither acts as cooperator and cheater and BY4741 is no longer cooperative in glucose or fructose media (non-public goods). Therefore, coevolution of both strains can be considered as evolution starting with isogenic population in non-public goods system.

To test this, three coevolution experiments were started with the BY4741 and ΔSUC2 (1:1) in 0.2% glucose, 0.2% fructose and mixture of 0.1% glucose and 0.1% fructose media independently. Three independent lines were propagated for each media. As neither of the strains has to cooperate and incur the fitness cost both are expected to have similar fitness values. Indeed, the initial ratio of frequency of cooperators to cheaters in all lines are ∼1:1 (Figure 3B).

Further the frequency of cooperators fluctuates across lines through different generations (**Figure 7A-C**). The adaptation trajectory is random in all the three lines. Either cheaters or cooperators evolved to do slightly better than its counterpart. This is expected given that the initial population is isogenic and that any of the two genotypes is capable of acquiring a beneficial mutation and increasing in frequency in any of the experiment’s environments. There was no significant increase in the frequency of either of the strains in the fructose and mixture of glucose and fructose media (p-value>0.3, unpaired t-test). In glucose media, there was a slight increase in the frequency of cooperators in two lines after 100 generations (p-value <0.05, unpaired t-test).

**Figure 7.**
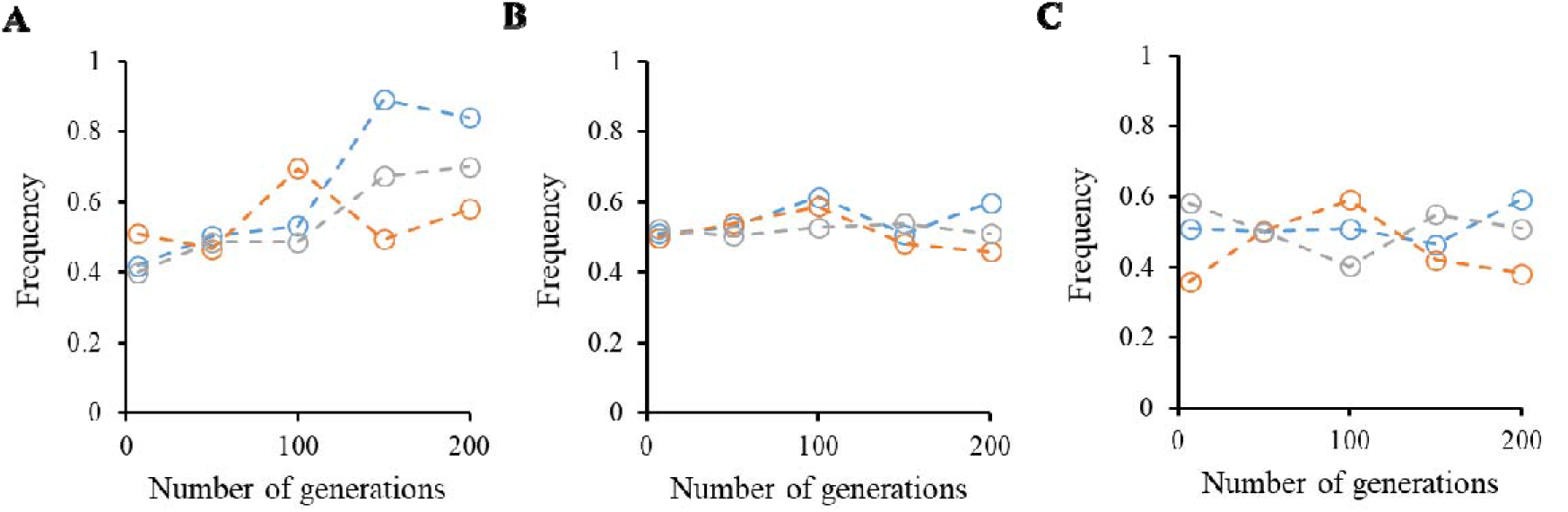
The coevolution of BY4741 and ΔSUC2 in different sugar medium was started with 1:1 ratio of both genotypes. Glucose and fructose utilization systems are not public good systems; hence both the strains can perform equally well. The ratio of frequency of cooperators to cheaters was quantified every 50 generations. The frequency of cooperators either increased or decreased every 50 generations in all the lines. **(A)** 0.2% glucose: There is a significant increase in the frequency of cooperators in the two of the lines after 200 generations compared to frequency at 7^th^ generation (p-value<0.05, unpaired t-test) **(B)** 0.1% glucose+0.1% fructose: There is no significant increase in the frequency of cooperators in any line after 200 generations compared to frequency at 7^th^ generation (p-value>0.8, unpaired t-test). **(C)** 0.2% fructose: There is no significant increase in the frequency of cooperators in any line after 200 generations compared to frequency at 7^th^ generation (p-value>0.9, unpaired t-test).

.The continuous increase in cooperator frequency was manifested only in sucrose. None of these non-public good systems showed the convergent adaptation of BY4741 to outperform ΔSUC2, which was found in sucrose. This implies the adaptation of cooperators to proliferate despite the cost was contingent on the coevolution in public good system such as sucrose.

### How do cooperators evolve in sucrose in the absence of cheaters?

The ancestor BY4741 was evolved for 200 generations in 0.2% sucrose medium in the absence of cheaters. This experiment was conducted to investigate whether cooperators, which evolve in sucrose without cheaters in the system, outperform the cheaters when ancestor cheaters are introduced into the system. After every 50th generation, the evolving cooperator and the ancestor ΔSUC2 were competed and the frequency of the two strains was quantified. Cooperators in two of the three lines were performing better than ancestor cheaters after 50 generations of evolution (**Figure 8A**). At 200^th^ generation, in the presence of ancestor cheater, the evolved cooperator of all 3 lines were performing significantly better than ancestor cooperator (p-value < 0.0006, paired t-test) [Figure 6A]. The competition assay was conducted with evolved cooperator at 200^th^ generation and ancestor cheater. The fraction of evolved cooperator was more than 50%. These results are not significantly different from what we observed in the competition between coevolved cooperator (200^th^ generation) and ancestor cheater (p-value = 0.16, unpaired t-test) (**Figure 8B, 5A)**. The growth assay was conducted with evolved cooperator BY4741 at 200th generation. The growth rate of cooperator had significantly increased in one line, while other two had the same growth rate or slightly greater than that of the ancestor BY4741 (**Figure 7C**). Thus, in a public-good system, the cooperator adapts to maintain cooperation. This is true for cases when the cooperator is growing in presence of cheaters, or by itself. In reality, the two cases are not quite distinct from each other since a single mutation in SNP can lead to evolution of cheaters in the case where cooperators are the only genotype present.

**Figure 8.**
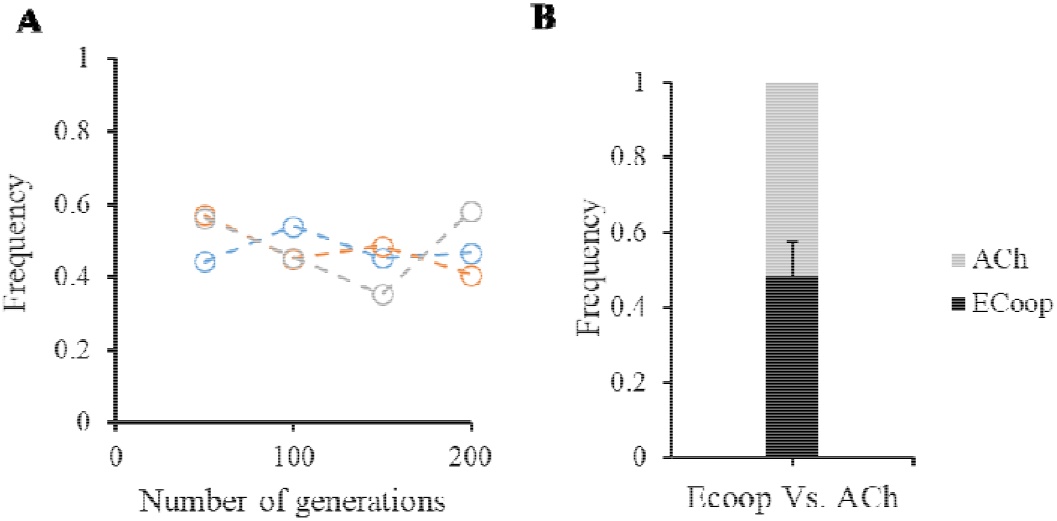
**(A)** The cooperator BY4741 was grown in 0.2% sucrose. Every 50 generations, competition assay was conducted with evolving cooperator and ancestor cheater to quantify their frequency values. At 200^th^ generation, in the presence of ancestor cheater, evolved cooperator of all 3 lines were performing better than ancestor cooperator (p-value < 0.0006 paired t-test). **(B)** Competition assay was conducted with evolved cooperator (Ecoop) at 200th generation with ancestor cheater (Ach). The frequency of evolved cooperators was ∼0.5. The ratio of frequency of cooperator to cheater does not vary significantly from the results observed in the competition between coevolved cooperator (200th generation) and ancestor cheater (p-value ∼ 0.16, unpaired t-test).

### The molecular bases of the cooperator adaptation in sucrose when coevolved with cheaters

The genome sequence data of ancestor BY4741 and cooperators coevolved with cheaters in 0.2% sucrose (coevolved BY4741) of all the six experimental lines were obtained by the whole genome sequencing. The data was further analysed to identify the mutations underlying the adaptation of cooperators to outperform cheaters. All mutations in the evolved lines are listed in the Table S1-S3.

Each of the six lines has a few non-synonymous single nucleotide variants (SNVs), as compared to the ancestor (**Table S1**). Coevolution line 1 had acquired single nucleotide variations in FLO9 gene. The gene FLO9 is involved in flocculation of cells. But BY4741 being a derivative of S288c, it has a nonsense mutation in FLO8 gene(33), a transcriptional activator of FLO family genes(34). BY4741 lacks flocculation as a result of the nonsense mutation. The coevolved cooperator of line 2 was found to have an ALD5 gene mutation. The mutant form of this gene, which is involved in the production of electron chain transport, has been found to have increased alcohol dehydrogenase activity and decreased respiration(35). In cooperator of line2, in addition to an SNV in the ALD5 gene, another mutation in the HRK1 gene was found. The protein kinase HRK1 controls the plasma membrane (H+)-ATPase. Expression of (H+)-ATPase is stimulated by glucose metabolism(36). Glucose metabolism causes an increase in growth rate by activating ATPase and elevating protein pump activity(37). An SNV mutation is observed in MYO1 gene of cooperator of line 5. MYO1 encodes the protein involved in cytokinesis(38). From this analysis the molecular basis of adaptation of the cooperators, when grown together with the cheater strains is not apparent.

However, we also note that one of the convergent changes observed in all the lines is duplication of regions of the chromosomes containing hexose transport sugars. In addition, in three of the lines there is a duplication of the region of the chromosome 9 which contains the gene SUC2 (**Table S2**).

## Discussion

Maintenance of cooperation between members of the same or individuals belonging to different species is an open problem in evolutionary biology. A number of mechanisms have been reported which explain this phenomenon(3-5, 9, 12, 18, 39, 40). In addition, theory has also helped provide insights towards possible mechanisms which explain maintenance of cooperation(17, 18, 20, 21, 41). In this work, we report an evolution experiment where we track increase in fitness of the cooperators, when grown together with cheater cells. The environment of growth makes the cooperators susceptible to be exploited by the cheaters. However, during the course of the evolution experiment, the cooperators increase in frequency at the cost of the cheaters.

Genome sequencing of the evolved co-operators suggests that this increase in fitness was accomplished via multiple mechanisms. The mutational targets (for SNPs) are wide-ranging between the six evolved lines, with little convergence.

However, we also note striking features of convergent evolution. All lines have regions of chromosomes duplicated which have hexose transporters. Three of the lines report convergent evolution, where region of chromosome 9 is duplicated. The SUC2 gene in this region of the chromosome, codes for the invertase which is responsible for hydrolysis of sucrose into its constituent monosaccharides, glucose and fructose.

At the start of the evolution experiment, we anticipated that increased privatization of the public good (SUC2) or the resulting monosaccharides could be possible mechanisms via which the cooperators might increase in fitness. In this context, we anticipated that the mutational targets might influence transport of the sugars across membrane. However, the sequencing results show that in half the lines, SUC2 is duplicated. We hypothesize that increased production of SUC2p leads to increased rate of hydrolysis, and therefore, doubling the amount of monosaccharides available to the cooperator. At low resource availability, this doubling could likely be critical in ensuring survival in a cheater population. This effect coupled with increase hexose transporter expression, likely leads to adaptation in a public good system, and helps the cooperator increase frequency in presence of cheaters.

An alternate but a strategy with a similar goal (of increasing availability of the monosaccharides) has been observed previously. The work by Koschwanez and team(42) reported that, individual cells were found to aggregate in small clusters, where each cell could benefit from the monosaccharide released form the immediate neighbors. If a cooperative phenomenon could link clustering, that would help spread of cooperation as a phenomenon in the population.

While the precise mechanism via which cooperation prevails in the population (in all lines of our experiment) is not known, it is apparent that cooperation as a phenomenon can survive via multiple mechanisms. Investigation of mechanistic details of how cooperation prevails in a population, via the mutations identified in this work, is currently under study.

## Materials and Methods

### Strains used

The two strains used in the study were BY4741 and ΔSUC2. The BY4741 is a S288c derivative(43). BY4741(genotype: *MATa his3*Δ*1 leu2*Δ*0 met15*Δ*0 ura3*Δ*0 SUC2*) has the SUC2 gene which encodes for the invertase. The ΔSUC2 is a cheater strain that has SUC2 gene knocked out from BY4741 and has a hygromycin-resistant (HPH) gene cassette instead.

To replace SUC2 gene with hygromycin-resistant (HPH) gene, the HPH gene cassette from plasmid pUG75 was amplified using the forward primer (5’- CAA GCA AAA CAA AAA GCT TTT CTT TTC ACT AAC GTA TAT GAT GCT TTT GCG CAG GTC GAC AAC CCT TAA T- 3’) and reverse primer (5’- TTT AGA ATG GCT TTT GAA AAA AAT AAA AAA GAC AAT AAG TTT TAT AAC CTA GTG GAT CTG ATA TCA CCT A -3’) which are homologous to the site of integration. The PCR products were transformed into BY4741 by electroporation, using Eppendorf eporator. The transformants where selected on YPD media containing Hygromycin (200 μg/mL). The knockout was confirmed using the forward primer (5’- CTC TTG TTC TTG TGC TTT TT -3’) and reverse primer (5’- ATT CTT TGA AAT CAT AAA GT -3’).

### Coevolution experiment

The two strains BY4741 and ΔSUC2 were revived from freezer stock on YPD (0.5% Yeast Extract, 1% Peptone and 2% Dextrose (w/v)). After 48 h of incubation, a single colony of BY4741 and ΔSUC2 from YPD plates were inoculated in a non-repressing and non-inducing glycerol-lactate media separately and incubated for 42-48 h. The coevolution experiment was conducted in four independent sugar environments: (a) sucrose, (b) glucose, (c) fructose and (d) mixture of glucose and fructose (CSM - 0.671% YNB with nitrogen base and 0.05% complete amino acid mixture; containing 0.2% of sugar). To begin with the evolution, 50 μL of each of the glycerol-lactate pregrown BY4741 (cooperator) and ΔSUC2 (cheater) culture were transferred in 1:1 ratio to a 5ml of CSM containing 0.2% sucrose (a public good system). Six independent replicates of coevolution were started in sucrose environment. Similarly, cheaters and cooperators were coevolved in other three sugar (non-public goods) media independently (0.2% glucose, 0.1% fructose, 0.1% glucose+0.1% fructose). Three replicate lines were initiated for each of the control environments.

In each of these lines, culture was incubated for 24 h at 30□ and 250 rpm. After 24 h, the culture was propagated by subculturing (1:100) to fresh media. A 1:100 dilution corresponds to roughly ∼6.7 generations per transfer. The experiment was continued till 200 generations. After every 50 generations, frequency of both the cooperator and cheater was quantified by using the fact that the cheater is resistant to Hygromycin. Freezer stocks of evolved BY4741 and ΔSUC2 strain were prepared after every 100 generations.

### Evolution experiment

To begin with the evolution of only cooperators, 50 μL of glycerol-lactate pre-grown BY4741 (cooperator) was transferred to a 5ml of CSM containing 0.2% sucrose. There were three replicates of evolution in sucrose environment. In each of these lines, culture was incubated for 24 h at 30□ and 250 rpm; cultures were propagated by subculturing (1:100) to a fresh sugar media after every 24 h. The experiment was conducted till 200 generations.

### To count the frequency of cooperators and cheaters

The coevolution cultures were diluted and plated on YPD plates to obtain ∼200 colonies. The colonies from YPD were transferred to YPD media containing Hygromycin (200 μg/mL) and incubated for 24 h. On this media, only the cheaters grew as BY4741 (cooperator) is sensitive to hygromycin and does not grow.

### Growth kinetics

The cells were revived from freezer stock on YPD plates. After 40-48 h of incubation, a single colony from the YPD plate was transferred to glycerol-lactate media and grown till saturation (∼48 h). The saturated culture was sub-cultured (1:100) to the media of interest to an initial OD of 0.05. Optical density (OD600) was measured using Thermoscientific Multiscan Go. Depending on the growth medium, the kinetics of growth were assessed every 4 to 6 hours until the culture reached saturation (6 h for sucrose and 4 h for glucose or fructose).

### Competition assays

The pair of strains to be competed were grown independently in glycerol-lactate for 48 h. Around 10^5^ cells from each of these glylac-lactate cultures were transferred to 5ml CSM containing sugar of interest with the concentrations used for coevolution experiments and grown for 24 h. The frequency of both the strains was calculated by above mentioned method.

### Whole-genome sequencing

The strain to be sequenced was revived from freezer stock on YPD plate. Single colony was inoculated and grown in 10mL liquid YPD for 10-15 h. The cells were harvested and genomic DNA was isolated following the *Saccharomyces cerevisiae* genomic DNA isolation protocol(44). The quantity and purity were measured using a Nano-spectrophotometer from Eppendorf (basic). High throughput sequencing was performed on Illumina platform with a read length of 150bp and minimum sequencing coverage of 100X. The sequencing data and the reads’ base qualities were stored in FASTQ file. The sequencing data is available at: https://www.ncbi.nlm.nih.gov/sra/PRJNA977882.

### Sequencing data analysis

The whole genome sequencing reads were stored in FASTQ files. These pairwise reads were aligned to the S288C (assembly: GCA_000146045.2 - R64) reference genome obtained from NCBI database. The reference-based sequence alignment was performed using BWA tool(version-0.7.17)(45). The raw alignment files were processed using Picard and Samtools (version-1.16.1)(46) to remove duplicates and sort the BAM files obtained from BWA tool. Genome analysis toolkit(GATK version - 4.3.0.0) was used to call the variants such as Single Nucleotide Variants (SNV) or Insertion-deletions (Indels) using haplotypecaller method(47). GATK-VariantFiltration was used to filter SNVs (SNPs) and Indels. SNPs and Indels were only taken into consideration if they passed the following filter: “QD < 2.0 || FS > 60.0 || MQ < 50.0 || HaplotypeScore > 13.0 || DP<50 || MappingQualityRankSum < -12.5 || ReadPosRankSum < -8.0”. Theses variants were annotated by Ensemble Variant Effect Predictor (VEP)(48).

Copy number variation (CNV) analysis: CNVpytor was used to call regions with copy number variations(49). By analysing the BAM files obtained by BWA tool, CNVpytor call regions with CNVs using a read-depth based approach. The bin size was set to 100bp. To select reliable CNVs, the following filter was used: e-val1<0.0001 (P-value calculated using i-test statistics between RD difference in the region and global (i.e., across whole genome) mean); q0<0.2 (fraction of reads mapped with zero quality within call region-to remove the reads mapped to multiple regions).

### Statistical analysis

All the p-values were obtained by performing unpaired student t-test to quantify the difference between ancestor and evolved strain unless mentioned otherwise.

## Acknowledgements

This work was funded by a DBT/Wellcome Trust (India Alliance) grant (Award No. IA/S/19/2/504632) to SS. NR is supported by the Prime Minister’s Research Fellowship (PMRF ID 1301163).

## References

1. Fritts RK, McCully AL, McKinlay JB. 2021. Extracellular Metabolism Sets the Table for Microbial Cross-Feeding. Microbiol Mol Biol Rev 85.

2. Butaite E, Baumgartner M, Wyder S, Kummerli R. 2017. Siderophore cheating and cheating resistance shape competition for iron in soil and freshwater Pseudomonas communities. Nat Commun 8:414.

3. Wingreen NS, Levin SA. 2006. Cooperation among microorganisms. PLoS Biol 4:e299.

4. West SA, Diggle SP, Buckling A, Gardner A, Griffins AS. 2007. The social lives of microbes. Annual Review of Ecology Evolution and Systematics 38:53–77.

5. Ozkaya O, Xavier KB, Dionisio F, Balbontin R. 2017. Maintenance of Microbial Cooperation Mediated by Public Goods in Single- and Multiple-Trait Scenarios. J Bacteriol 199.

6. Lerch BA, Smith DA, Koffel T, Bagby SC, Abbott KC. 2022. How public can public goods be? Environmental context shapes the evolutionary ecology of partially private goods. PLoS Comput Biol 18:e1010666.

7. Sandoz KM, Mitzimberg SM, Schuster M. 2007. Social cheating in Pseudomonas aeruginosa quorum sensing. Proc Natl Acad Sci U S A 104:15876–81.

8. Damore JA, Gore J. 2012. Understanding microbial cooperation. J Theor Biol 299:31–41.

9. Waite AJ, Shou W. 2012. Adaptation to a new environment allows cooperators to purge cheaters stochastically. Proc Natl Acad Sci U S A 109:19079–86.

10. Sanchez A, Gore J. 2013. feedback between population and evolutionary dynamics determines the fate of social microbial populations. PLoS Biol 11:e1001547.

11. Miller MB, Bassler BL. 2001. Quorum sensing in bacteria. Annu Rev Microbiol 55:165–99.

12. Wang M, Schaefer AL, Dandekar AA, Greenberg EP. 2015. Quorum sensing and policing of Pseudomonas aeruginosa social cheaters. Proc Natl Acad Sci U S A 112:2187–91.

13. Wechsler T, Kummerli R, Dobay A. 2019. Understanding policing as a mechanism of cheater control in cooperating bacteria. J Evol Biol 32:412–424.

14. Travisano M, Velicer GJ. 2004. Strategies of microbial cheater control. Trends Microbiol 12:72–8.

15. Santorelli LA, Kuspa A, Shaulsky G, Queller DC, Strassmann JE. 2013. A new social gene in Dictyostelium discoideum, chtB. BMC Evol Biol 13:4.

16. Yan H, Asfahl KL, Li N, Sun F, Xiao J, Shen D, Dandekar AA, Wang M. 2019. Conditional quorum-sensing induction of a cyanide-insensitive terminal oxidase stabilizes cooperating populations of Pseudomonas aeruginosa. Nat Commun 10:4999.

17. Gore J, Youk H, van Oudenaarden A. 2009. Snowdrift game dynamics and facultative cheating in yeast. Nature 459:253–6.

18. MaClean RC, Fuentes-Hernandez A, Greig D, Hurst LD, Gudelj I. 2010. A mixture of “cheats” and “co-operators” can enable maximal group benefit. PLoS Biol 8.

19. Greig D, Travisano M. 2004. The Prisoner’s Dilemma and polymorphism in yeast SUC genes. Proc Biol Sci 271 Suppl 3:S25–6.

20. Prajapat MK, Shroff I, Brajesh RG, Saini S. 2016. Analysis of a strategy for cooperating cells to survive the presence of cheaters. Mol Biosyst 12:3338–3346.

21. Rauch J, Kondev J, Sanchez A. 2017. Cooperators trade off ecological resilience and evolutionary stability in public goods games. J R Soc Interface 14.

22. De La Fuente G, Sols A. 1962. Transport of sugars in yeasts. II. Mechanisms of utilization of disaccharides and related glycosides. Biochim Biophys Acta 56:49–62.

23. Lagunas R. 1993. Sugar transport in Saccharomyces cerevisiae. FEMS Microbiol Rev 10:229–42.

24. Neigeborn L, Carlson M. 1984. Genes affecting the regulation of SUC2 gene expression by glucose repression in Saccharomyces cerevisiae. Genetics 108:845–58.

25. Winge O, Roberts C. 1952. The relation between the polymeric genes for maltose raffinose, and sucrose fermentation in yeasts. Cr Trav Lab Carlsberg Ser Physiol 25:141–71.

26. Hawthorne DC, Mortimer RK. 1960. Chromosome Mapping in Saccharomyces: Centromere-Linked Genes. Genetics 45:1085–110.

27. Tkacz JS, Lampen JO. 1973. Surface distributon of invertase on growing Saccharomyces cells. J Bacteriol 113:1073–5.

28. Ozcan S, Vallier LG, Flick JS, Carlson M, Johnston M. 1997. Expression of the SUC2 gene of Saccharomyces cerevisiae is induced by low levels of glucose. Yeast 13:127–37.

29. Weinhandl K, Winkler M, Glieder A, Camattari A. 2014. Carbon source dependent promoters in yeasts. Microb Cell Fact 13:5.

30. Lutfiyya LL, Johnston M. 1996. Two zinc-finger-containing repressors are responsible for glucose repression of SUC2 expression. Mol Cell Biol 16:4790–7.

31. Chen A, Sanchez A, Dai L, Gore J. 2014. Dynamics of a producer-freeloader ecosystem on the brink of collapse. Nat Commun 5:3713.

32. Craig Maclean R, Brandon C. 2008. Stable public goods cooperation and dynamic social interactions in yeast. J Evol Biol 21:1836–43.

33. Liu H, Styles CA, Fink GR. 1996. Saccharomyces cerevisiae S288C has a mutation in FLO8, a gene required for filamentous growth. Genetics 144:967–78.

34. Kobayashi O, Suda H, Ohtani T, Sone H. 1996. Molecular cloning and analysis of the dominant flocculation gene FLO8 from Saccharomyces cerevisiae. Mol Gen Genet 251:707–15.

35. Kurita O, Nishida Y. 1999. Involvement of mitochondrial aldehyde dehydrogenase ALD5 in maintenance of the mitochondrial electron transport chain in Saccharomyces cerevisiae. FEMS Microbiol Lett 181:281–7.

36. Goossens A, de La Fuente N, Forment J, Serrano R, Portillo F. 2000. Regulation of yeast H(+)-ATPase by protein kinases belonging to a family dedicated to activation of plasma membrane transporters. Mol Cell Biol 20:7654–61.

37. Serrano R. 1983. In vivo glucose activation of the yeast plasma membrane ATPase. FEBS Lett 156:11–4.

38. Watts FZ, Shiels G, Orr E. 1987. The yeast MYO1 gene encoding a myosin-like protein required for cell division. EMBO J 6:3499–505.

39. Allen B, Nowak MA. 2013. Cooperation and the fate of microbial societies. PLoS Biol 11:e1001549.

40. O’Brien S, Lujan AM, Paterson S, Cant MA, Buckling A. 2017. Adaptation to public goods cheats in Pseudomonas aeruginosa. Proc Biol Sci 284.

41. Traulsen A, Nowak MA. 2006. Evolution of cooperation by multilevel selection. Proc Natl Acad Sci U S A 103:10952–5.

42. Koschwanez JH, Foster KR, Murray AW. 2011. Sucrose utilization in budding yeast as a model for the origin of undifferentiated multicellularity. PLoS Biol 9:e1001122.

43. Brachmann CB, Davies A, Cost GJ, Caputo E, Li J, Hieter P, Boeke JD. 1998. Designer deletion strains derived from Saccharomyces cerevisiae S288C: a useful set of strains and plasmids for PCR-mediated gene disruption and other applications. Yeast 14:115–32.

44. Dymond JS. 2013. Preparation of genomic DNA from Saccharomyces cerevisiae. Methods Enzymol 529:153–60.

45. Li H, Durbin R. 2009. Fast and accurate short read alignment with Burrows-Wheeler transform. Bioinformatics 25:1754–60.

46. Li H, Handsaker B, Wysoker A, Fennell T, Ruan J, Homer N, Marth G, Abecasis G, Durbin R, Genome Project Data Processing S. 2009. The Sequence Alignment/Map format and SAMtools. Bioinformatics 25:2078–9.

47. McKenna A, Hanna M, Banks E, Sivachenko A, Cibulskis K, Kernytsky A, Garimella K, Altshuler D, Gabriel S, Daly M, DePristo MA. 2010. The Genome Analysis Toolkit: a MapReduce framework for analyzing next-generation DNA sequencing data. Genome Res 20:1297–303.

48. McLaren W, Gil L, Hunt SE, Riat HS, Ritchie GR, Thormann A, Flicek P, Cunningham F. 2016. The Ensembl Variant Effect Predictor. Genome Biol 17:122.

49. Suvakov M, Panda A, Diesh C, Holmes I, Abyzov A. 2021. CNVpytor: a tool for copy number variation detection and analysis from read depth and allele imbalance in whole-genome sequencing. Gigascience 10.

