## Supplement Tables for "Maintenance of cooperation in a yeast population in a public-good driven system"

**Table S1. Single nucleotide polymorphism (SNPs) found in all evolution lines.**

| Line | Chrmosome | Gene | cDNA pos: SNP | Amino acid change |
| --- | --- | --- | --- | --- |
| 1 | Chr I: 27044 | FLO9 | 934 C-->T | 309V->I |
|  | Chr I: 27035 | FLO9 | 925 T-->C | 312I->V |
|  | Chr VII: 859401 | UBR1 | T->C | downstream_gene_variant |
|  | Chr VII:859401 | TIM13 | T->C | downstream_gene_variant |
| 2 | Chr V: 304849 | ALD5 | 820 A->G | 274 I->V |
|  | Chr XV: 824858 | HRK1 | 10 G->T | 4 L->I |
|  | Chr XV:516943 | OST2 | C- > A | upstream_gene_variant |
|  | Chr V: 115662 | URA3 | C- > A | upstream_gene_variant |
|  | Chr XV :516943 | PIN2 | C- > A | upstream_gene_variant |
|  | Chr XV :516943 | RAS1 | C- > A | downstream_gene_variant |
| 3 | Chr XIII :384585 | STV1 | 1283 G->A | 428 R->K |
| 4 | Chr X: 715095 | DAN4 | 646 G->A | 216 P->S |
|  | Chr XV: 118032 | NDJ1 | A->C | upstream_gene_variant |
| 5 | Chr VIII: 156178 | MYO1 | 4513 G->A | 1505 A->T |
|  | Chr XIV: 175245 | RAD50 | G- > A | upstream_gene_variant |
|  | Chr XII :412652 | SLX4 | G- > A | upstream_gene_variant |
|  | Chr XII : 412650 | SLX4 | C -> T | upstream_gene_variant |
|  | Chr XIV: 175245 | NRD1 | G- > A | upstream_gene_variant |
| 6 | Chr XVI: 799541 | CTF4 | 308 T->G | 103 I->S |
|  | Chr XV: 118032 | NDJ1 | A->C | upstream_gene_variant |
|  | Chr XVI: 799541 | MSS18 | T->G | downstream_gene_variant |

**Table S2. List of putative chromosomal locations with copy number variations.**

| **Line** | **Chromosome** | **Region: start** | **Region: end** | **Genes in the region** |
| --- | --- | --- | --- | --- |
| 1 | Chr I | 163901 | 204600 | BUD14 |
|  | Chr III | 293601 | 316700 | GIT1 |
|  | Chr VI | 1 | 86300 | MOB2, RIM15 |
|  | Chr VI | 225501 | 270200 | HXK1 |
|  | Chr VII | 1 | 78000 | HXK2, RTG2 |
|  | Chr XIV | 14501 | 98700 | HXT14 |
|  | Chr XIV | 700901 | 784400 | HXT17 |
|  | Chr XV | 1 | 29100 | IMA2, HXT11 |
| 2 | Chr VI | 250201 | 270200 | HXK1 |
|  | Chr IX | 389401 | 439900 | FLO11 |
|  | Chr XV | 1 | 28200 | IMA2, HXT11 |
| 3 | Chr I | 164001 | 189400 | BUD14 |
|  | Chr III | 293401 | 316700 | GIT1 |
|  | Chr VII | 1 | 20700 | HXK1 |
|  | Chr XIV | 723001 | 784400 | HXT17 |
|  | Chr XV | 1 | 31200 | IMA2, HXT11 |
| 4 | Chr I | 25701 | 140800 | FLO9, CDC24, CDC19, MYO4, FLC2 |
|  | Chr I | 163901 | 204600 | BUD14 |
|  | Chr IV | 1 | 98800 | HXT15 |
|  | Chr IV | 1437201 | 1532000 | GIN4 |
|  | Chr V | 1 | 115900 | BUD16 |
|  | Chr VI | 214301 | 270200 | HXK1 |
|  | Chr VII | 1 | 78800 | HXK2, RTG2 |
|  | Chr VII | 1028601 | 1091000 | IMA1, MAL13, MAL12 |
|  | Chr IX | 23401 | 48500 | SUC2 |
|  | Chr XIV | 14501 | 103300 | HXT14 |
|  | Chr XIV | 684701 | 784400 | HXT17 |
|  | Chr XV | 45801 | 94800 | MSN1 |
| 5 | Chr I | 1 | 140900 | SEO1, PAU8, FLO9, TDA8 |
|  | Chr I | 164001 | 204600 | BUD14 |
|  | Chr V | 53501 | 115900 | BUD16 |
|  | Chr VII | 1 | 78400 | HXK2, RTG2 |
|  | Chr VII | 1013201 | 1091000 | IMA1, MAL13, MAL12 |
|  | Chr IX | 23401 | 53900 | SUC2 |
|  | Chr XIV | 14401 | 100700 | HXT14 |
|  | Chr XIV | 708801 | 784400 | HXT17 |
|  | Chr XV | 1 | 62200 | HXT11, IMA2 |
|  | Chr XV | 997601 | 1081500 | PFK27 |
| 6 | Chr I | 26701 | 140800 | FLO9, CDC24, CDC19, MYO4, FLC2 |
|  | Chr I | 164001 | 204600 | BUD14 |
|  | Chr VI | 197901 | 270200 | HXK1 |
|  | Chr VII | 1 | 72700 | HXK2, RTG2 |
|  | Chr VII | 1033301 | 1091000 | IMA1, MAL13, MAL12 |
|  | Chr IX | 23401 | 52300 | SUC2 |
|  | Chr X | 601801 | 639600 | BUD4 |
|  | Chr XIII | 795901 | 924500 | DIA1 |
|  | Chr XIV | 14501 | 100700 | HXT14 |
|  | Chr XIV | 702401 | 784400 | HXT17 |
|  | Chr XV | 1 | 118300 | IMA2, HXT11 |
|  | Chr XV | 969401 | 1081400 | PFK27 |
